# Integrative Multiomic Analyses of Diabetic Neuropathic Pain in Dorsal Root Ganglia: Proteomics, Phospho-proteomics, and Metabolomics

**DOI:** 10.1101/2022.07.31.502193

**Authors:** Megan Doty, Sijung Yun, Yan Wang, Minghan Hu, Margaret Cassidy, Bradford Hall, Ashok B. Kulkarni

## Abstract

Diabetic peripheral neuropathy (DPN) is characterized by spontaneous pain in the extremities. Incidence of DPN continues to rise with the global diabetes epidemic. However, there remains a lack of safe, effective analgesics to control this chronic painful condition. Dorsal root ganglia (DRG) contain soma of sensory neurons and modulate sensory signal transduction into the central nervous system. In this study, we aimed to gain a deeper understanding of changes in molecular pathways in the DRG of DPN patients with chronic pain. We recently reported transcriptomic changes in the DRG with DPN. Here, we expand upon those results with integrated metabolomic, proteomic, and phospho-proteomic analyses to compare the molecular profiles of DRG from DPN donors and DRG from control donors without diabetes or chronic pain. Our analyses identified decreases of select amino acids and phospholipid metabolites in the DRG from DPN donors, which are important for cellular maintenance. Additionally, our analyses revealed changes suggestive of extracellular matrix (ECM) remodeling and altered mRNA processing. These results reveal new insights into changes in the molecular profiles associated with DPN.

## Introduction

Chronic pain is a growing health burden, impacting an estimated 20% of adults in the United States [1]. Among chronic pain cases, an estimated 25% can be attributed to neuropathic pain [2]. A common painful neuropathic condition is Diabetic Peripheral Neuropathy (DPN), which is caused by Diabetes Mellitus (DM). Type II Diabetes Mellitus (T2D) is an epidemic that affects 462 million people globally, a number expected to grow markedly, with 50% of patients developing DPN [3, 4, 5]. DPN is defined as a length-dependent sensorimotor polyneuropathy. DPN is associated with hyperglycemic pathology as well as hyperlipidemia; however, a mechanism of DPN development remains unclear [5, 6, 7]. DPN typically manifests in a ‘stocking and glove’ pattern in which patients’ lower limbs are most affected, consistent with disease of long sensory neurons. Additionally, DPN is associated with axonal demyelination and degeneration leading to nerve dysfunction and possible cell death [8]. Exponential growth in the diabetes epidemic and a lack of an effective curative treatment for DPN make it increasingly imperative to gain insights into the mechanisms of disease progression [9, 2, 10].

Dorsal root ganglia (DRG) contain the cell bodies of peripheral sensory neurons that relay pain form the periphery to the central nervous system [11]. It is well established that transcriptomic changes occur in DRG in response to injury and in association with neuropathic pain development [11]. Our previously published mRNA sequencing (RNA-seq) study suggested an association between DPN, increased expression of immune-related genes, and decreased expression of neuronal genes in the human DRG [12]. However, due to translational and post-translational regulation, expression of a specific mRNA is not a perfect indicator of the abundance of the corresponding protein, especially in the context of neurological dysfunction [13–18]. Despite this, to the best of our knowledge, no study to date has explored translational and post-translational regulation in human DRG. Protein abundance and post-translational modifications (PTM) are central to cellular function, but easily deregulated in response to physiological changes [19]. Phosphorylation is an important PTM upon which many cellular processes are reliant. Additionally, metabolites can be influenced directly by endogenous and exogenous factors and are direct players in biochemical processes, thus offering a strong descriptor of molecular phenotype. A few studies have identified metabolomic and proteomic trends suggestive of disrupted energy metabolism and mitochondrial function in DRG of diabetic rodents [20–23]. Better understanding of molecular regulation with painful conditions in human systems is vital for development of novel therapeutics [24].

Therefore, we aimed to integrate previous transcriptomic findings [12] with metabolomic, proteomic, and phospho-proteomic data to offer a comprehensive understanding of the biochemical changes associated with DPN in human DRG. Presently, we profiled small molecular weight metabolites and lipids, as well as proteins and phospho-peptides in human DRG samples from DPN donors and non-diabetic controls. We identified decreased amino acids and phospholipid metabolites, as well as alterations in abundance of extracellular matrix (ECM) proteins and phosphorylation of RNA binding proteins. From this, we suggest a role of disrupted amino acid and protein metabolism in the development of DPN.

## Results

### Tissue donor cohort characteristics

For metabolomic analysis DRG were obtained from 7 DPN donors and 10 non-diabetic control donors. For proteomic and phospho-proteomic analyses, DRG were obtained from 5 DPN donors and 5 non-diabetic control donors. Within both cohorts of donors, no significant differences were observed between donor groups in age, sex, or BMI (Table 1). A significant difference was observed in donors on ACE inhibitors in the metabolomic cohort, but no ACE inhibitor metabolites were detected in our analysis [25]. A non-significant trend was observed in differences in age in our metabolomic cohort, this was addressed in our statistical analysis of metabolite data.

**Table 1:** Donor characteristics for DRG tissues used in metabolomic and proteomic analyses

|  | Metabolomic Cohort |  |  | Proteomic Cohort |  |  |
| --- | --- | --- | --- | --- | --- | --- |
|  | Control, N=10 | DPN, N = 7 | p-value | Control, N=4 | DPN, N = 5 | p-value |
| **DRG Storage** |  |  | 0.9<sup>1</sup> |  |  |  |
| Snap Frozen | 5 (50%) | 4 (57%) |  | 4 (100%) | 4 (100%) |  |
| RNAlater | 5 (50%) | 3 (43%) |  |  |  |  |
| **Sex** |  |  | 0.9<sup>1</sup> |  |  | 0.5<sup>1</sup> |
| Female | 5 (50%) | 3 (43%) |  | 3 (75%) | 2 (40%) |  |
| Male | 5 (50%) | 4 (57%) |  | 1 (25%) | 3 (60%) |  |
| **Age** | 46.8 (7.6) | 53.4 (9.8) | 0.1<sup>2</sup> | 51.8 (2.5) | 53.4 (9.8) | 0.7<sup>2</sup> |
| **BMI** | 27.4 (6.1) | 30.6 (8.6) | 0.7<sup>2</sup> | 35.0 (11.4) | 30.6 (8.6) | 0.5<sup>2</sup> |
| **Race/Ethnicity** |  |  | 0.5<sup>1</sup> |  |  | 0.5<sup>1</sup> |
| African American | 2 (20%) | 3 (43%) |  | 1 (25%) | 2 (40%) |  |
| Hispanic/Latino | 2 (20%) | 3 (43%) |  | 0 (0%) | 1 (20%) |  |
| White | 6 (60%) | 2 (29%) |  | 3 (75%) | 2 (40%) |  |
| **Cause of Death** |  |  | 0.6<sup>1</sup> |  |  | 0.6<sup>1</sup> |
| Anoxia/Cardiovascular | 2 (20%) | 3 (43%) |  | 1 (25%) | 2 (40%) |  |
| CVA/ICH/Stroke | 4 (40%) | 4 (43%) |  | 1 (25%) | 2 (40%) |  |
| MVA/Head Trauma/Blunt Injury | 4 (40%) | 1 (14%) |  | 2 (50%) | 2 (40%) |  |
| **Medications** |  |  |  |  |  |  |
| Vasodilators | 3 (30%) | 1 (14%) | 0.6<sup>1</sup> | 3 (60%) | 2 (40%) | 0.9<sup>1</sup> |
| Opioids | 1 (10%) | 3 (43%) | 0.2<sup>1</sup> | 0 (0%) | 2 (40%) | 0.4<sup>1</sup> |
| Gabapentin | 0 (0%) | 2 (29%) | 0.2<sup>1</sup> | 0 (0%) | 2 (40%) | 0.4<sup>1</sup> |
| Insulin | 0 (0%) | 2 (29%) | 0.2<sup>1</sup> | 0 (0%) | 3 (60%) | 0.2<sup>1</sup> |
| Metformin | 0 (0%) | 2 (29%) | 0.2<sup>1</sup> | 0 (0%) | 2 (40%) | 0.2<sup>1</sup> |
| Ca<sup>2+</sup> Channel Blockers | 1 (10%) | 4 (57%) | 0.1<sup>1</sup> | 0 (0%) | 2 (40%) | 0.4<sup>1</sup> |
| $\beta$ Blockers | 1 (10%) | 4 (57%) | 0.1<sup>1</sup> | 0 (0%) | 3 (60%) | 0.2<sup>1</sup> |
| ACE Inhibitors | 1 (10%) | 5 (71%) | 0.035<sup>1*</sup> | 0 (0%) | 3 (60%) | 0.2<sup>1</sup> |
| Statins | 1 (10%) | 4 (57%) | 0.1<sup>1</sup> | 1 (20%) | 2 (40%) | 0.9<sup>1</sup> |
| **Comorbidities** |  |  |  |  |  |  |
| Cardiac Disease | 1 (10%) | 4 (57%) | 0.1<sup>1</sup> | 1 (20%) | 2 (40%) | 0.9<sup>1</sup> |
| Hypertension | 4 (40%) | 6 (86%) | 0.13<sup>1</sup> | 3 (60%) | 4 (80%) | 0.9<sup>1</sup> |
| Neurological Disease | 2 (20%) | 1 (14%) | 0.058<sup>1</sup> | 1 (20%) | 3 (60%) | 0.5<sup>1</sup> |
| Cancer | 0 (0%) | 1 (14%) | 0.4<sup>1</sup> | 0 (0%) | 0 (0%) |  |
| Diabetes | 0 (0%) | 7 (100%) | 5e-05<sup>1*</sup> | 0 (0%) | 5 (100%) | 0.008<sup>1*</sup> |

Results given as N (%) or mean (standard deviation)
<sup>1</sup>Fisher's Exact Test
<sup>2</sup> T-test

Post-mortem tissue donation networks offer the best opportunity for analyses of human DRGs. L4, L5, and S1 DRGs were obtained as these ganglia contain the primary afferent neurons that innervate the distal extremities of the foot where patients with DPN often experience pain (Supplemental Figure S1a). Donor medical history confirms neuropathy for 10 years or more in the DPN cohort as observed from the available timeline. However, with current limitations in obtaining human DRG [26], we were unable to obtain detailed clinical measures of pain hypersensitivity in DRG donors. We have earlier reported pathological evaluation of DRG tissues from the DPN and control donors showing variation in pathology in both donor groups [12]. As reported earlier [12], within the non-diabetic control group, pathology ranges from apparently normal tissue to moderate ganglionic cell loss, whereas within the DPN group, pathology ranges from apparently normal tissue to moderate to severe ganglionic cell loss.

For our present study, profiles of protein, DNA, and RNA degradation products as well as indicators of cellular damage were evaluated to characterize tissue quality (Supplementary Figure S1b). Abundance of these degradation products would be abnormally high if tissues were poorly handled or stored. We do note that one sample has exceptionally high levels of hypoxanthine, inosine, and certain amino acids (Supplementary Figure S1b). A second sample is noted to have high levels of certain amino acids as well (Supplementary Figure S1b). However, for both samples, these increases were not consistent across all amino acids. Additionally, diphosphates ADP and GDP were particularly low in these two samples. As such, we determined no DRG samples with abnormal degradation were included in our analysis.

### Metabolomic profiling suggests disrupted amino acid and phospholipid metabolism

Metabolomic profiling was performed with 7 DRG from donors with DPN and 10 DRG from non-diabetic controls (Table 1). Capillary electrophoresis mass spectrometry (CE-MS) and liquid chromatography mass spectrometry (LC-MS) collectively identified 327 metabolites in donor DRG tissues. 165 metabolites were detected in more than 80% of samples and included in further analysis (Figure 1a).

**Figure 1.**
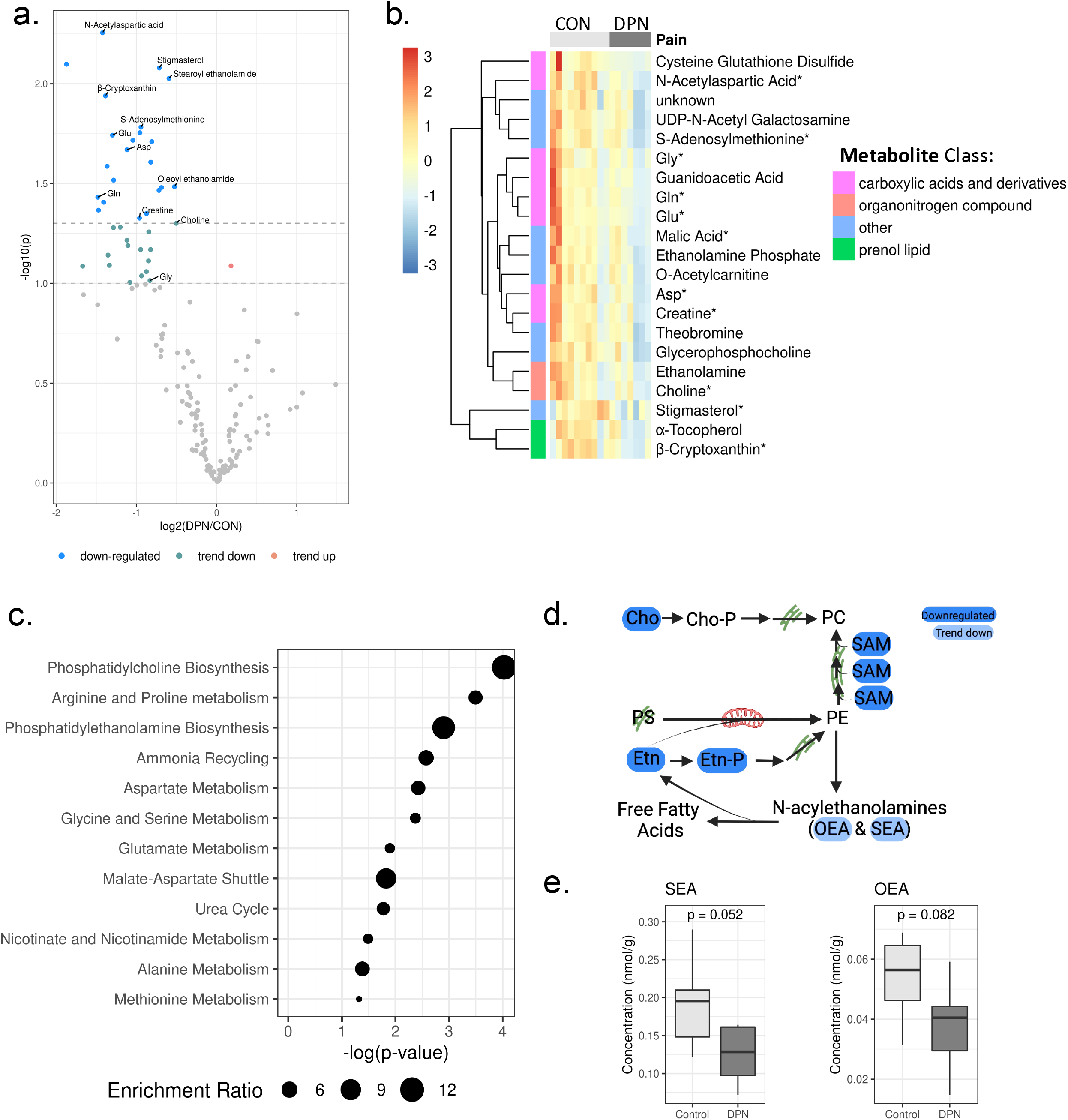
Decreased amino acids and phospholipid metabolites with DPN. A) Volcano plot showing metabolite abundance according to ratio of DPN/control and p-values before covariate control. Significantly regulated metabolites considered to be those with p-value < 0.05. Metabolites with trends in regulation identified with p-value < 0.1. Dashed lines represent p-values 0.1 and 0.05 before covariate control. B) Heat map showing scaled and centered relative abundance of differentially regulated metabolites (p-value after covariate control < 0.05). Asterisks (*) indicates metabolites which were validated with quantitative panels. Row-wise annotations indicate metabolite class, metabolite classes represented by only one metabolite are grouped together as other. CON indicates control samples, DPN indicates diabetic peripheral neuropathy. C) Metabolite enrichment analysis with SMPDB library shows enrichment in phospholipid biosynthesis and amino acid metabolism. Annotations shown are significant with p-value <0.05. Enrichment ratio is observed number of metabolites divided by expected number metabolites in metabolite set. D) Schematic representation of PC and PE biosynthesis pathways as well as derivation of N-acylethanolamines, created with BioRender. E) Box plot showing trends in decreased PE derived lipids, SEA and OEA. Indicated p-value was calculated after covariate control.

Dimensionality reduction with principal component analysis (PCA) suggested the influence of donor pain condition on metabolite profiles to be confounded by variables such as sample storage condition (Supplementary Figure S2). Linear regression models were applied to candidate metabolites to account for variation due to sample storage condition, donor age, and donor sex. 21 metabolites were identified as significantly altered as a result of neuropathic pain (p-value adjusted for covariates < 0.05) after this covariate adjustment (Figure 1b, Supplementary Table S1). All differentially regulated metabolites were decreased in the DRG from donors with DPN.

Of the 21 identified metabolites, 8 are classified as carboxylic acids and derivatives (N-acetylaspartic acid (NAA), creatine (Cr), glycine (Gly), aspartic acid (Asp), glutamine (Gln), glutamate (Glu), guanidoacetic acid, and cysteine glutathione disulfide) (Figure 1b). We noted additional non-significant decreases in carboxylic acids and derivatives including asparagine (ratio (DPN/control) = 0.46, p-value adjusted for covariates = 0.057) and ornithine (ratio (DPN/control) = 0.38, p-value adjusted for covariates = 0.058) (Supplementary Table S1). Similarly, our enrichment analysis revealed significantly enriched metabolite sets related to arginine and proline metabolism, aspartate metabolism, glycine and serine metabolism, and glutamate metabolism (Figure 1c). Because these observations suggest alterations in amino acids, the concentrations of proteinogenic amino acids were then examined with quantitative panels (Supplementary Figure S3). Three differentially regulated amino acids (Gln, Gly, and Glu) were observed to be among the highest abundance proteinogenic amino acids in the DRG (Supplementary Figure S3). Of note, non-proteogenic amino acid NAA, was the most significantly decreased metabolite (ratio (DPN/control) = 0.37, p-value adjusted for covariates = 0.00995). Given that NAA is highly abundant in the nervous system and neuronally important [27], regulation of NAA was confirmed with a quantitative panel (Supplementary Table S1).

Among our significantly enriched metabolite sets, we also observed an enrichment for biosynthetic pathways of two phospholipids, phosphatidylcholine (PC) and phosphatidylethanolamine (PE) (Figure 1c). Organonitrogen compounds choline (Cho, ratio (DPN/control) = 0.71, p-value adjusted for covariates = 0.049, Figure 1b) and ethanolamine (Etn, ratio (DPN/control) = 0.48, p-value adjusted for covariates = 0.0068, Figure 1b) are involved in both pathways (Figure 1d). Ethanolamine phosphate (Etn-P, ratio (DPN/control) = 0.56, p-value adjusted for covariates = 0.027, Figure 1b) is also involved in both pathways (Figure 1d), whereas S-adenoylmethionine (SAM) (ratio (DPN/control) = 0.52, p-value adjusted for covariates = 0.0164, Figure 1b) is involved in PC synthesis (Figure 1d). To further support altered phospholipid dynamics, we observed trending losses in two N-acylethanolamines, oleoyl-ethanolamide (OEA, ratio (DPN/control) = 0.69, p-value adjusted for covariates = 0.082, Figure 1e) and stearoyl-ethanolamide (SEA, ratio (DPN/control) = 0.66, p-value adjusted for covariates = 0.052, Figure 1e). Given that PE is the sole endogenous source of OEA and SEA, these metabolites were selected for validation. Quantitative panels confirmed the identity and trend in differential abundance of OEA and SEA (Figure 1e, Supplementary Table S1), as well as differential regulation of Cho and SAM (Supplementary Table S1).

### Proteomic profile shows extracellular matrix remodeling

Proteomic profiling was performed with 5 DRG derived from donors with DPN and 5 DRG derived from control donors without diabetes or chronic pain (Table 1). After sample prep, total peptide abundance in 1 non-diabetic control sample was abnormally low, this sample was excluded from further analysis. 6186 proteins were identified in human DRG samples. 247 proteins were identified as significantly differentially regulated (Benjamini-Hochberg adjusted p-value < 0.05, Figure 2a). Dimensionality reduction analyses with PCA and t-distributed Stochastic Neighbor Embedding (t-SNE) did not show clear sample grouping by neuropathic pain (Supplementary Figure S4). We believe this was due to some influence of covariate variables and as such used linear modeling to adjust for age and sex. 41 proteins were identified as significantly altered after covariate control (p-value adjusted for covariates < 0.05, Supplementary Table S2), of these 5 are regulated by fold change greater than 1.5 (|log2(DPN/control)| > 0.58, Figure 2b).

**Figure 2.**
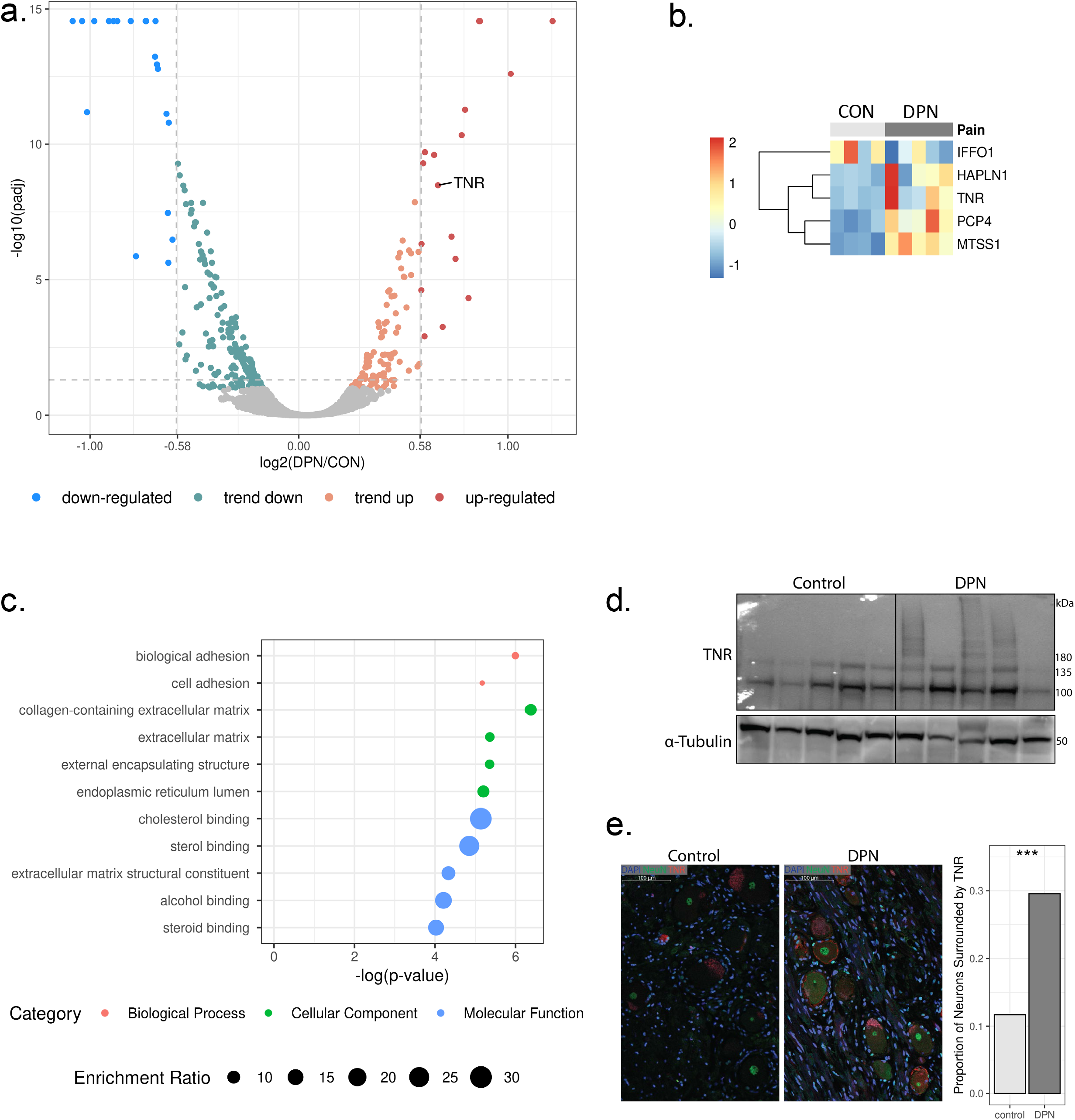
Extracellular matrix remodeling with DPN. A) Volcano plot showing protein abundance according to ratio of DPN/control and adjusted p-values calculated with ProteomeDiscoverer. Vertical lines at 0.58 and −0.58 represent an abundance ratio of 1.5 and 0.67 respectively. Proteins are considered up regulated or downregulated with adjusted p-value < 0.05 and fold change > 1.5. Proteins are considered trending up or down with 0.05 < p-value < 0.1 or p-value < 0.05 and fold change < 1.5. B) Heat map showing relative abundance of differentially regulated proteins, identified after covariate control (p-value after covariate control < 0.05, abundance ratio > 1.5 or < 0.67). CON indicates control samples, DPN indicates diabetic peripheral neuropathy. C) Results of enrichment analysis with differentially regulated proteins (p-value after covariate control < 0.05). Top gene ontology annotations include biological adhesion and extracellular matrix. D) Western blot shows heavy glycosylated TNR products only for DPN samples. E) IHC results showing TNR (red) localization surrounding neurons (NeuN, green) in DRG sections. DAPI is labeled in blue. A z-test compared proportion of neurons surrounded by TNR in control and DPN derived DRG sections (p-value = 7.6e-11).

From statistical overrepresentation test using Panther, differentially regulated proteins (p-value adjusted for covariates < 0.05) showed enrichment in gene ontology annotations including ECM and biological adhesion (Figure 2c). Changes to the ECM and cellular adhesion is highlighted by upregulation of the neuron-specific protein, tenascin R (TNR, ratio (DPN/control) = 1.59, p-value adjusted for covariates = 0.048). Using an alternative method, we sought to further explore regulation of ECM proteins as identified by mass spec. TNR is 150 kDa protein and subject to PTMs such as glycosylation which increase its molecular weight. Western blot analysis of TNR revealed truncated TNR protein products (<150 kDa) in all DRG samples (Figure 2d). Heavy TNR protein products (>150kDa) were detected in 3 DPN samples and 0 control samples (Figure 2d). This reveals a TNR regulation pattern unique to samples within the DPN group.

Considering TNR is an ECM protein, we aimed to further explore regulation of this protein with visualization of its localization in the DRG. To do so, we performed immunohistochemistry with DRG tissue sections. NeuN was used to label neurons in the DRG. TNR fluorescence was observed surrounding select neurons in the DRG (Figure 2e). We quantified the proportion of neurons in the DRG that are surround by TNR. In control DRG sections, of 453 neurons evaluated, 53 were surrounded by TNR. In DPN DRG sections, of 433 neurons evaluated, 128 neurons were surrounded by TNR. In total, 11.7% of neurons in DRG derived from control donors were surrounded by TNR, whereas 29.6% of neurons in DRG derived from the DPN donors were surrounded by TNR (p-value = 7.6e-11, Figure 2e).

### Phospho-proteomic results show changes to mRNA processing

Using the sample samples as in proteomic profiling, samples were enriched for phospho-peptides and subjected to profiling by mass spectrometry. 7842 phosphorylated peptides mapping to 2533 master proteins were identified. Of these, 153 phospho-peptides were identified as differentially regulated between DPN and non-diabetic control groups (Benjamini-Hochberg adjusted p-value < 0.05, Figure 3a). Dimensionality reduction analyses with PCA and t-SNE again did not show clear separation of samples by neuropathic pain (Supplementary Figure S6). We again assumed an influence of covariates and we used linear regression to adjust for age and sex. 27 phospho-peptides were identified after covariate control (p-value adjusted for covariates < 0.05, Supplementary Table S3), 20 of these were regulated by a fold change greater than 1.5 (|log2(DPN/control)| > 0.58, Figure 3b).

**Figure 3.**
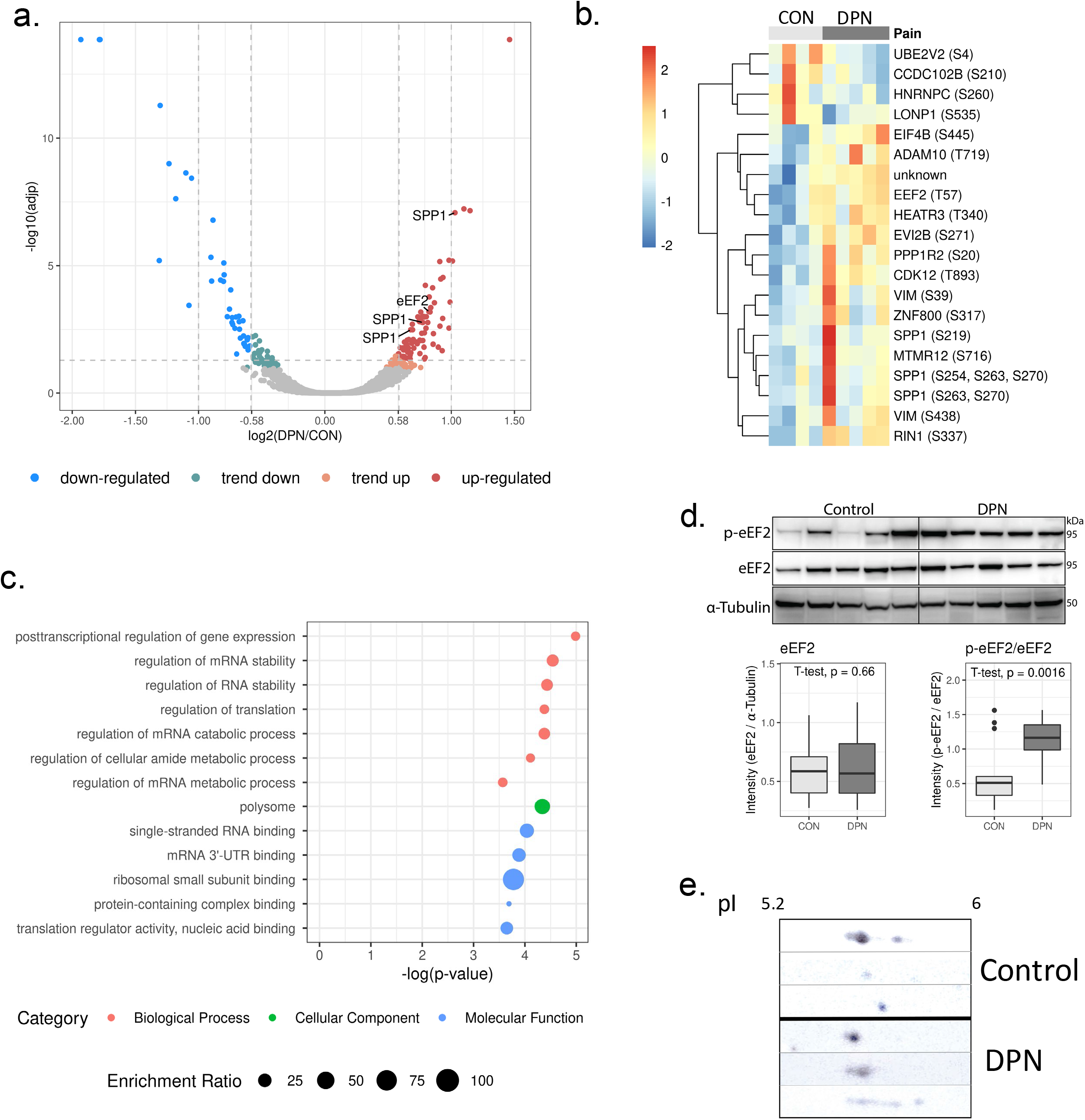
Disrupted protein translation in DPN. A) Volcano plot showing phospho-peptide abundance according to ratio of DPN/control and adjusted p-values calculated in ProteomeDiscoverer. Vertical lines at 0.58 and −0.58 represent 1.5-fold change. Vertical lines at 1 and −1 represent an abundance ratio of 2 or 0.5 respectively. Phospho-peptides are considered up regulated or downregulated with adjusted p-value < 0.05 and fold change > 1.5. Phospho-peptides are considered trending up or down with 0.05 < p-value < 0.1 or p-value < 0.05 and fold change < 1.5. B) Heat map showing relative abundance of differentially regulated phospho-peptides (p-value after covariate control < 0.05, abundance ratio > 1.5 or < 0.67). CON indicates control samples, DPN indicates diabetic peripheral neuropathy. C) Results of enrichment analysis with differentially regulated phospho-proteins (p-value after covariate control < 0.05). Top gene ontology annotations include regulation of mRNA stability and regulation of translation. D) Western blot results confirmed increased phosphorylation of eEF2 with DPN. E) 2D gel showing different migration patterns of SPP1 in DPN and control samples, isoelectric point (pI) is noted at the top. Within DPN samples, SPP1 is observed to have acidic, phosphorylated residues, whereas SPP1 in control samples has a more basic migration pattern.

Differential regulation of phospho-peptides could be explained by either differential phosphorylation or differential regulation of master proteins abundance. To explore this, we looked for overlap in proteomic and phospho-proteomic data. We found that not only are SPP1 phospho-peptides increased, but SPP1 protein is also slightly increased (ratio (DPN/control) = 1.32, p-value after covariate control = 0.012, Supplementary Table S3). However, abundance ratios of SPP1 phospho-peptides show larger fold changes, demonstrating phosphorylation differences. All other identified phospho-proteins were either not detected, or not significantly regulated in background proteome data (Supplementary Table S3). This confirms that identified differentially regulated phospho-proteins are in fact differentially phosphorylated.

3 phospho-peptides derived from SPP1 were upregulated by abundance ratio greater than 1.5 (pS219, ratio (DPN/control) = 1.6, p-value after covariate control = 0.017; pS263 and pS270, ratio (DPN/control) = 2.0, p-value after covariate control = 0.012; pS254 and pS263 and pS270, ratio (DPN/control) = 1.7, p-value after covariate control = 0.036, Figure 3b). Additional phospho-peptides derived from SPP1 were also increased, however with a smaller fold change (pS375 and pS280, ratio (DPN/control) = 1.5, p-value after covariate control = 0.012; pS303, ratio (DPN/control) = 1.4, p-value after covariate control = 0.0218; pS254, ratio (DPN/control) = 1.478, p-value after covariate control = 0.02, Supplementary Table S3). Due to phosphorylation changes at multiple sites, we hypothesized that overall phosphorylation state and therefore isoelectric point of SPP1 is altered with DPN. As expected, 2D gel electrophoresis confirmed differential migration patterns of SPP1 in DPN and control groups, with more acidic migration of SPP1 in DPN samples (Figure 3e). As phosphorylation is an acidic PTM, this supports increased phosphorylation of SPP1 in the DRG with DPN.

Using a statistical overrepresentation test in Panther, enrichment analysis of differentially regulated phospho-peptides revealed enrichment in annotations involving post-transcriptional regulation of gene expression and regulation of mRNA stability (Figure 3c). Central to this is eukaryotic translation elongation factor (eEF2), which shows increased phosphorylation at the T57 residue (ratio (DPN/control) = 1.77, p-value adjusted for covariates = 0.046, Figure 3b). Because this phospho-site is well studied and known to inhibit translation activity of eEF2 [28, 29], we selected this phospho-protein for further study. We used western blot to validate mass spectrometry findings. Western blot analysis confirmed increased phosphorylation of eEF2 at the T57 residue in DRGs from DPN donors (ratio (DPN/control) = 1.78, p-value = 0.0016, Figure 3d).

### Multi-omics integration suggests contribution of disrupted amino acid and protein metabolism in neuronal dysfunction

Using an IPA, we looked for commonalities in disease and function annotations among our omics datasets. Synthesis of protein, metabolism of protein, and synthesis of amino acids were suggested to be decreased, while uptake of amino acids was possibly identified as increased (Table 2). As expected, these annotations are drawn from decreases in amino acids (Table 2). Phospho-proteins eEF2 and SPP1 are also related to decreased synthesis and metabolism of protein (Table 2). Additionally, SPP1, eEF2, TNR, certain amino acids (NAA, Glu), and certain phospholipid metabolites (SAM, Cho) are annotated as contributing to increased progressive neurological disorder (Table 2). Altogether, this links amino acid and protein metabolism with neurological dysfunction in DPN (Supplementary Figure S6b). This also links changes to ECM proteins and phospholipid metabolism to neurological dysfunction in DPN (Supplementary Figure S6b).

**Table 2:**
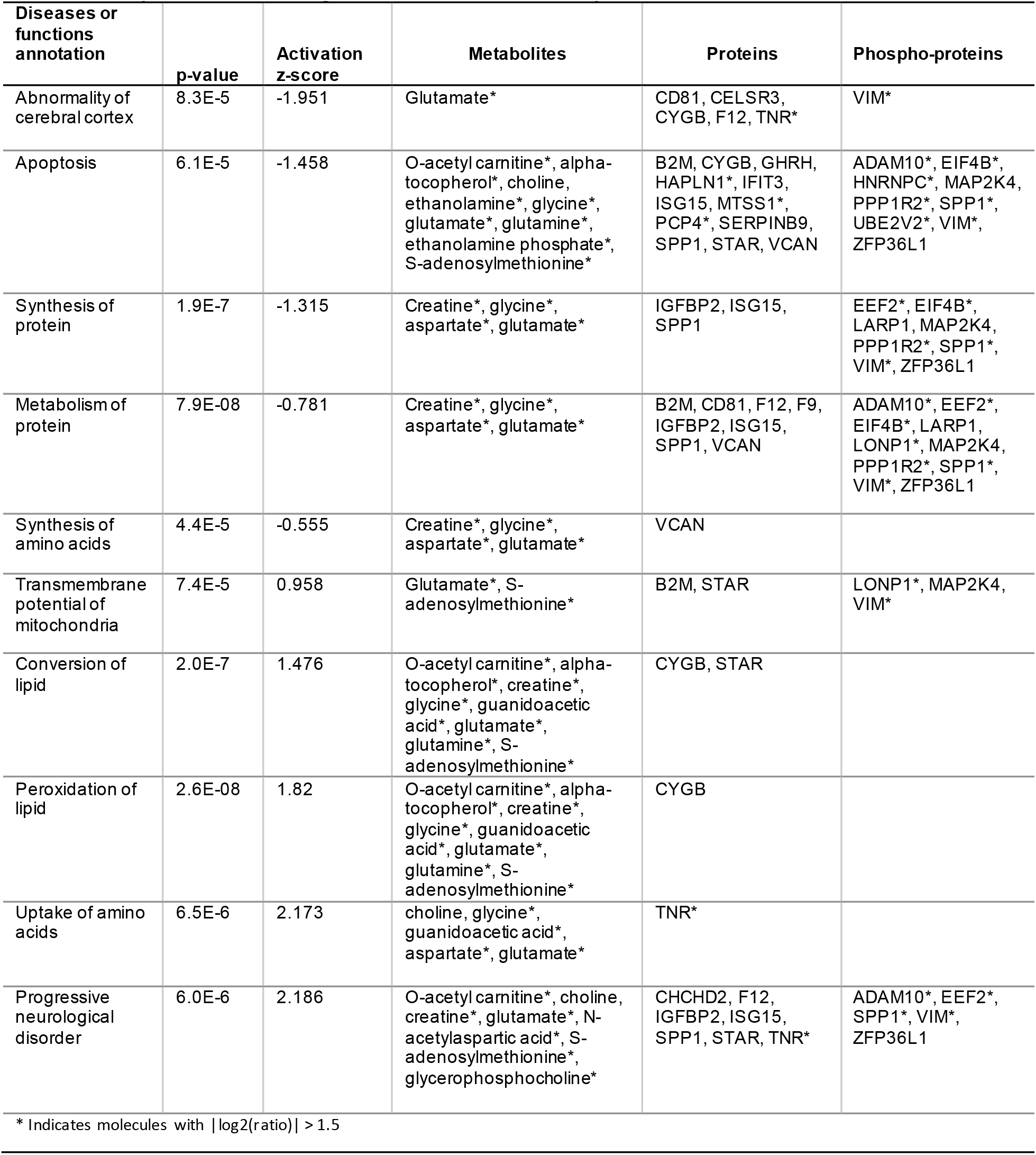
Top results of integrative enrichment analysis with IPA.

## Discussion

DPN is a common form of chronic pain for which there is no effective curative treatment. Demyelination, axonal degeneration, and neuronal dysfunction and death are linked to progression of pain in DPN [8]. Here, we used a multi-omics approach to gain a comprehensive understanding of molecular alterations associated with this pathology. Given the difficulties in obtaining human DRG samples, this descriptive omics-based study reveals unique information necessary for further elucidating the etiology and progression of DPN. Our metabolomic study revealed depletion of select amino acids in the DRG in our DPN cohort (Figure 1b, c) as well as decreased phospholipid-related metabolites (Figure 1c, d). In proteomic and phospho-proteomic data we find changes to structural ECM proteins (Figure 2b, c) and differential phosphorylation of RNA binding proteins (Figure 3b, c) in the DRG with DPN. Increased abundance of TNR (Figure 2d) as well as hyperphosphorylation of eEF2 (Figure 3d) and SPP1 (Figure 3e) in DPN were confirmed using western blot. Integration of this data using IPA suggests decreased amino acids and altered phosphorylation of RNA binding proteins contribute to altered protein synthesis and neurological disorder in the DRG in DPN. (Table 2, Supplementary Figure S6a, b).

In our metabolomics analysis, we record decreases in 8 carboxylic acid and derivatives, including proteogenic amino acids Glu, Gln, Gly, and Asp (Figure 1b). Of note, IPA reveals a link between amino acids Glu and NAA, and progressive neurological disorders (Table 2, Supplementary Figure S6a, b). Although homeostasis for most amino acids is maintained by transport processes, intracellular Glu and Asp levels are influenced most by energy metabolism [30] (Supplementary Figure S6b). Observations of decreased Gly and Gln are consistent with alterations in plasma levels of diabetic patients [31, 32]. Such states of amino acid depletion have been linked to cell death and neurodegeneration [33–35]. However, whether amino acid depletion in the DRG is sufficient to produce degeneration observed in DPN is unclear. Additionally, Gly can function as an inhibitory neurotransmitter; however, a causal relation between decreased Gly in diabetes and neuronal hyperexcitability in DPN has not been studied [31].

Moreover, the non-proteinogenic amino acid NAA is decreased in our data (Figure 1b). NAA is synthesized in neuronal mitochondria by the enzyme NAT8L [36] (Supplementary Figure S6a). Interestingly, the gene transcript for the enzyme was decreased in RNA-seq data [12]; however this same change was note detected in our proteomic data. This discrepancy could be related to post-transcriptional regulation or sampling effects. Nonetheless, decreased NAA is important in the peripheral nervous system as it is utilized by Schwann cells to synthesize myelin [37] (Supplementary Figure S6a). Neuronal injury and degeneration decrease NAA, suggesting the differential regulation of NAA in DRG is specific to the neuropathic phenotype in DPN [27, 38]. NAA’s role in myelin synthesis also provides a possible mechanism for demyelination in DPN (Supplementary Figure S6a, b).

We also report decreases in metabolites linked to phospholipid biosynthesis (Figure 1c, d, e). Specifically, differentially regulated metabolites are metabolically related to PC and PE (Figure 1c). We also see a trend with DPN towards decreased SEA, a PE derived compound that has been reported to have an analgesic effect [39]. Phospholipids are also important in mitochondrial function and myelin maintenance [40, 41] (Supplementary Figure S6b). PC is a high abundance phospholipid in peripheral myelin [41]. The short half-life of this molecule [41] suggests a link between PC biosynthesis and demyelination in DPN (Supplementary Figure S6b).

Among significantly altered proteins, we observe enrichment of proteins in the ECM (Figure 2c). The ECM has been implicated in painful conditions by regulating the neuronal microenvironment, influencing synaptic plasticity, and modulating cell signaling [42–44]. We observe increased TNR with DPN, especially surrounding neurons in the DRG (Figure 2d, e). TNR is also noted to be related to amino acid uptake and progressive neurological disorder in IPA (Table 2, Supplementary Figure S6b). Specifically, TNR has been reported to indirectly regulate Glu uptake [45]. A direct relation between amino acid depletion and TNR upregulation in the DRG is not clear. However, increased TNR does inhibit axonal regeneration [46], thus, implicating this protein in neuronal degeneration observed in the DRG with DPN.

Among differentially regulated phospho-proteins, we report enrichment in annotations relating to RNA binding and translation (Figure 3c). Changes in phosphorylation of RNA binding proteins is highlighted by hyperphosphorylation of eEF2 at the T57 residue (Figure 3b, c, Supplementary Table S3). Phosphorylation at this site is well known to inhibit protein translation [29] (Supplementary Figure S6b). Interestingly, eEF2 hyperphosphorylation can occur in response to ER stress [47], a phenomenon suggested by our previous transcriptomic analysis [12]. Further evidence of ER stress can be noted by the ER’s role in phospholipid biosynthesis [48] and the evidence of decreased phospholipid biosynthesis in our data (Figure 1c, 1d, Supplementary Figure S6c). ER stress is well known to contribute to chronic pain and neurodegeneration [49–51]. With the ER being a single continuous organelle, spanning the entity of the axon, long sensory neurons are particularly vulnerable to ER stress [52]. However, the myelinating activities of glial cells also make them susceptible to ER stress [53]. Further work is needed to explore cell-type specific ER-stress in DPN.

We also report hyperphosphorylation of SPP1 (Figure 3b, e). We note a slight increase in total SPP1; however, this result was not validated, and the fold change was minimal (ratio (DPN/control) = 1.32), Supplementary Table S2). SPP1 is marker for proprioceptors in the DRG and is linked to nerve injury and mechanical pain [54, 55]. SPP1 has also been reported to be linked to ER stress [56, 57], inflammation in diabetes and neurodegeneration [58, 59], and adhesion in the extracellular matrix [60] (Supplementary Figure S6b). However, these reports looked at total SPP1, less is known about the function of phosphorylated SPP1.

We previously reported a pro-inflammatory signature in the DRG with DPN [12]. TNR, which is increased in our data (Figure 2b, d), is tightly linked to neuroinflammation [61]. SPP1 is involved in inflammatory processes as well [58–60, 62] although the role of SPP1 phosphorylation in these mechanisms is not known. We also find minor changes in interferon induced protein with tetratricopeptide repeats 3 (IFIT3, ratio = 1.25, p-value after covariate control = 0.024, Supplementary Table S2) and interferon stimulated gene 15 (ISG15, ratio = 1.44, p-value after covariate control = 0.017, Supplementary Table S2), both of which play a role in immune signaling. Inflammation is known to contribute to progression of both T2D and DPN [63, 64]. Lack of an identification of a stronger inflammatory signature in proteomic data is likely a result of detection limits of our methods.

This study is limited by a small sample size brought on by the lack of available human DRG tissue in both disease and healthy conditions. Additionally, a lack of available clinical pain assessment of DRG donors limits the power of this study. A study with the resources to overcome these limitations has yet to be powered and would allow valuable information to build off these preliminary results [26]. Out of an abundance of caution, we used strict covariate controls, however this may have added to limitations in our statistical power. Although determined to be minimal, postmortem changes likely limited the power of this study, as did detection limits of mass spectrometry. Despite this, our study offers an important preliminary look into metabolite and protein dynamics in a disease state in the human DRG. Further work is needed to explore this regulation in larger cohort of donors. Additionally, future studies are needed to explore mechanistic contributions of identified molecules in the development of pain in DPN.

In this pilot study, we explored for the first time, metabolite, protein, and phospho-protein regulation in the human DRG. Furthermore, comparison of non-diabetic and DPN derived DRG suggests a role of disrupted amino acid and protein metabolism in neuronal dysfunction in DPN. We propose use of findings here to advise further studies with DPN. In depth understanding of molecular regulation identified here could offer novel insights into the etiology and development of DPN.

## Methods

### Sample Acquisition

DRG used in this study were acquired from the cadaveric donors with informed consent of the next of kin (Anabios, San Diego, CA). We obtained approval for carrying out these studies from the Office of Human Subjects Research Protection (OHSRP), National Institutes of Health, Bethesda, MD, USA. L4, L5 and S1 DRGs were collected under cold ischemic conditions, within 3 hours of aorta cross-clamp. Donor medical history was obtained by trained interviewers from donor family members to the best of their knowledge. Despite lack of clinical data, organ donation offers unique and valuable insight molecular regulation in the soma of sensory neurons impacted in DPN.

### Metabolomic Analysis

#### Data Acquisition

S1 DRG samples (30-50 mg) were used for metabolite analysis (Human Metabolome Technologies-America, Boston, MA). For polar metabolites, frozen tissues were homogenized in acetonitrile in water (50%) with internal standards (20 μM). The supernatant was filtered through a 5 kDa filter, centrifugally concentrated, and resuspended in ultrapure water (50 μl). CE-TOFMS was carried out using Agilent CE-TOFMS system (Agilent Technologies Inc, Waldbronn, Germany) and fused silica capillary (50 μm × 80 cm) with HMT electrophoresis buffer and HMT sheath liquid. Capillary electrophoresis was run at 30 kV. The spectrometer scanned from mass/charge (m/z) 50 to 1,000.

For non-polar metabolites, frozen tissue was homogenized by vortexing with zirconium beads in 1% formic acid in acetonitrile containing internal standards (10 μM) and centrifuged (2,300 x g, 4°C, 5min). The supernatant was collected. The pellet was homogenized in 1% formic acid in acetonitrile and MilliQ-water (167 ul) followed by centrifugation (2,300 x g, 4°C, 5min). Supernatants were combined and filtered through a 3 kDa filter and a far filtered through phospholipid affinity column (Hybrid SPE phospholipid 55261-U, Supelco, Bellefonte, PA, USA). Filtrate was desiccated and resuspended in isopropanol (50%) in Mili-Q water. LC-TOFMS was carried out with Agilent 1200 series RRLC system SL (Agilent Technologies Inc, Wadbronn, Germany) with ODS column (2 x 50 mm, 2 μm) coupled to Agilent LC/MSD TOF MS system (Agilent Technologies Inc, Wadbronn, Germany). For chromatographic separation, mobile phase A was 0.1% HCOOH In H_2_O. Mobile Phase B was 0.1% HCOOH and 2 mM HCOONH_4_ in 65:30:5 isopropanol: acetonitrile: H_2_O. The gradient condition was 1% mobile phase B for 0.5 min, followed by a 13-minute ramp from 1% mobile phase B to 100% mobile phase B, followed by 100% mobile phase B for 6.5 min. Flow rate was 0.3 mL/min, column temperature was 40°C, injection volume was 1 μL, MS capillary voltage was 4 kV and 3.5 kV in ESI positive and ESI negative mode respectively, nebulizer pressure was 40 psi, gas flow was 10 L/min, and gas temperature was 350°C. The spectrometer scanned from mass/charge (m/z) 100 to 1700.

CE-TOFMS and LC-TOFMS data was processed with MasterHands (v2.17.1.11) for identification of peaks and quantification based on internal standards. Metabolites detected in fewer than 80% of samples were excluded from downstream analysis. Remaining missing values were imputed using half minimum method [65]. Identity and concentration of select metabolites was validated using external unlabeled standards. Human Metabolome Database was used to identify metabolite class and directly related enzymes.

### Proteomics

#### Sample Prep

Tissues were prepared for proteomic analysis using EasyPep Mini MS Sample Prep Kit according to manufacturer’s instructions (Thermo Fisher Scientific). Protein concentration was measured using Pierce BCA Protein Assay (Thermo Fisher Scientific). 100 μg aliquots of protein from each sample were reduced, alkylated, and digested. Remaining protein extracts were saved for electrophoresis and western blot. Samples were labeled with TMTpro 16-plex (Thermo Fisher Scientific) according to the manufacturer’s instructions. 10 μg of each sample was combined, fractionated using high pH fractionation kit (Thermo Fisher Scientific), and used for full proteome analysis. The remaining 90 μg of each sample was pooled for phospho-peptide enrichment with PTMScan Phospho-Enrichment IMAC FE-NTA Magnetic Beads (Cell Signaling Technology) according to the manufacturer’s instructions. The flow through was then enriched according to the High-Select SMOAC protocol (Thermo Scientific Scientific). In the SMOAC protocol, samples were sequentially enriched with High-Select TiO_2_ Phosphopeptide Enrichment Kit (Thermo Fisher Scientific) and High-Select Fe-NTA Phosphopeptide Enrichment Kit (Thermo Fisher Scientific).

#### Data Acquisition

All fractions were analyzed with nano LC-MS/MS with Thermo Scientific Fusion Lumos Tribrid mass spectrometer interfaced to a UltiMate3000 RSLCnano HPLC system (Thermo Fisher Scientific, San Jose, CA). For each analysis, 1 ug of corresponding fraction was loaded and desalted in an Acclaim PepMap 100 trap column (75 µm ’ 2 cm) at 4 μl/min for 5 min. Peptides were then eluted into a 75 μm× 250mm Accalaim PepMap 100 column (3 μm, 100 Å) and chromatographically separated using a binary solvent system consisting of A: 0.1% formic acid and B: 0.1% formic acid and 80% acetonitrile, at a flow rate of 300 nl/min. A gradient was run from 1% B to 42% B over 150 min, followed by a 5-min wash step with 80% B and a 10-min equilibration at 1% B before the next sample was injected. Precursor masses were detected in the Orbitrap at R=120,000 (m/z 200). HCD fragment masses were detected in the orbitrap at R=50,000 (m/z 200). Data-dependent MS/MS was carried out with top of speed setting, cycle time 2 s with dynamic exclusion of 20 s.

Proteome Discoverer Software (v2.5, Thermo Fisher Scientific, San Jose, CA) processed mass spectrometry data. MS spectra were searched against Homo sapiens and contaminant databases using the SEQUEST HT with PhosphoRS node for verification of phosphorylation sites. The following search parameters were used: enzyme: trypsin; maximum missed cleavage sites: 2; precursor mass tolerance: 10 ppm; fragment mass tolerance: 0.02 Da; dynamic modifications: oxidation (M), phosphorylation (S, T, Y), acetylation (protein N-terminus); static modifications: TMTpro (peptide N-terminus), carbamidomethyl), TMTpro (K); percolator strict FDR: <0.01, percolator relaxed FDR < 0.05.

For quantification, samples were normalized by total peptide abundance. Spectra with >50% isolation interference were excluded. Protein quantification was performed using unique and razor peptides, protein ratio calculation was protein abundance based, missing data was imputed using low abundance resampling. Proteome Discoverer reported background-based t-tests, with the Benjamini Hochberg correction.

### 2-D gel electrophoresis

Two-dimensional electrophoresis was performed with 500 ug of protein by Kendrick Labs, Inc. (Madison, WI) according to the carrier ampholine method of isoelectric focusing [66]. Isoelectric focusing was carried out in a glass tube of inner diameter 3.3 mm using 2.0% pH 3-10 isodalt Servalytes (Serva, Heidelberg, Germany) for 20,000 volt-hrs. Tube gels were then sealed to the top of a stacking gel that overlaid a 10% acrylamide slab gel (1.0 mm) and SDS slab gel electrophoresis was carried out. The gel was then transblotted onto PVDF membrane.

### Western Blotting

Proteins were separated with 4 to 12% bis-tris gels for eEF2 or 7% tris-acetate gel for TNR and transferred to a PVDF membrane. Membranes were blocked and incubated overnight with primary antibodies against pT57-eEF2 (1/1000, 2331, Cell Signaling Technologies), eEF2 (1/1000, 2332, Cell Signaling Technologies), TNR (1/2000, AF3865, R&D systems), alpha-tubulin (1/10,000, ab7291, Abcam), or SPP1 (1/1000, ab8448, Abcam). Membranes were washed and incubated with HRP conjugated secondary antibodies against rabbit IgG (1/10,000, 711-035-152, Jackson Immunoresearch), goat IgG (1/10,000, 705-035-003, Jackson Immunoresearch), or mouse IgG (1/10,000, ab6789, Abcam). Chemiluminescence was detected using ECL substrate and imaged with FlourChem M (Biotechne) or Amersham Imager 600 (GE Healthcare Life Sciences) for membranes from 2D gel. For detection of total eEF2 after detection of p-eEF2, the membrane was stripped with Restore PLUS Western Blot Stripping Buffer (Thermo Fisher Scientific).

### Immunohistochemistry

Formalin fixed paraffin embedded DRG tissue sections were deparaffinized and subject to citrate buffered antigen retrieval. Sections were stained using a primary antibody raised against TNR (1/50, AF3865, R&D systems) and secondary antibody conjugated to Rhodamine Red-X raised against goat IgG (1/100, 705-296-147, Jackson Immunoresearch). Sections were also stained with antibody raised against NeuN and conjugated to Alexa Flour488 (1/100, MAB377X, Sigma Aldrich). Sections were mounted using DAPI fluoromount-G (17984-24, Electron Microscopy Sciences). Fluorescence was imaged using a Nikon A1R HD25 Spectral microscope. Representative images at 10x magnification were used for analysis, images for figures were acquired at 20x magnification. The proportion of NeuN expressing cells which are also surrounded by TNR was compared between groups.

### Statistical Analysis

Statistical analyses were performed in R, unless otherwise specified. A T-test was used to assess statistical differences in metabolomic and western blot data, a z-test was used to assess statistical differences in immunohistochemistry data. For omics datasets, the R package stats (v3.6.2) was used to apply linear regression models to candidate molecules for covariate control. Enrichment analyses were performed with molecules with p-value after covariate control < 0.05. Metabolite enrichment was performed using metaboanalyst with Small Molecule Pathway Database (SMPDB). Proteome and phospho-proteome enrichment analysis was performed separately with Panther with p-values resulting from fisher’s exact test. Ingenuity Pathway Analysis (IPA) software (Ingenuity Systems, Mountain View, CA) was used for integrated enrichment of candidate metabolites proteins, and phospho-proteins.

## Acknowledgements

We would like to thank Anabios Corporation for procuring the human DRGs. We also thank Alexander Buko of Human Metabolome Technologies, America for acquisition and discussion of metabolomic data. We thank Dr. Pavan Auluck of NIMH for pathological evaluation of DRG tissues. We thank Dr. Bill Swaim and Dr. Duy Tran of the NIDCR imaging core for advisement with immunohistochemistry and imaging. We also thank Dr. Ranganath Muniyappa of NIDDK for his insightful comments and advisement. This research was supported by the Division of Intramural Research, NIDCR, NIH (ZIA-DE-000664-24, ZIA-DE-000751).

## Author Contributions Statement

MD, BH, and ABK conceptualized the project. MD, SY, YW carried out experiments and analyzed data. MD wrote original draft. MD, SY, YW, BH, MH, MC, and ABK reviewed and edited manuscript.

## Data Availability

Data is available upon request.

---

**Supplementary Figure S1.**
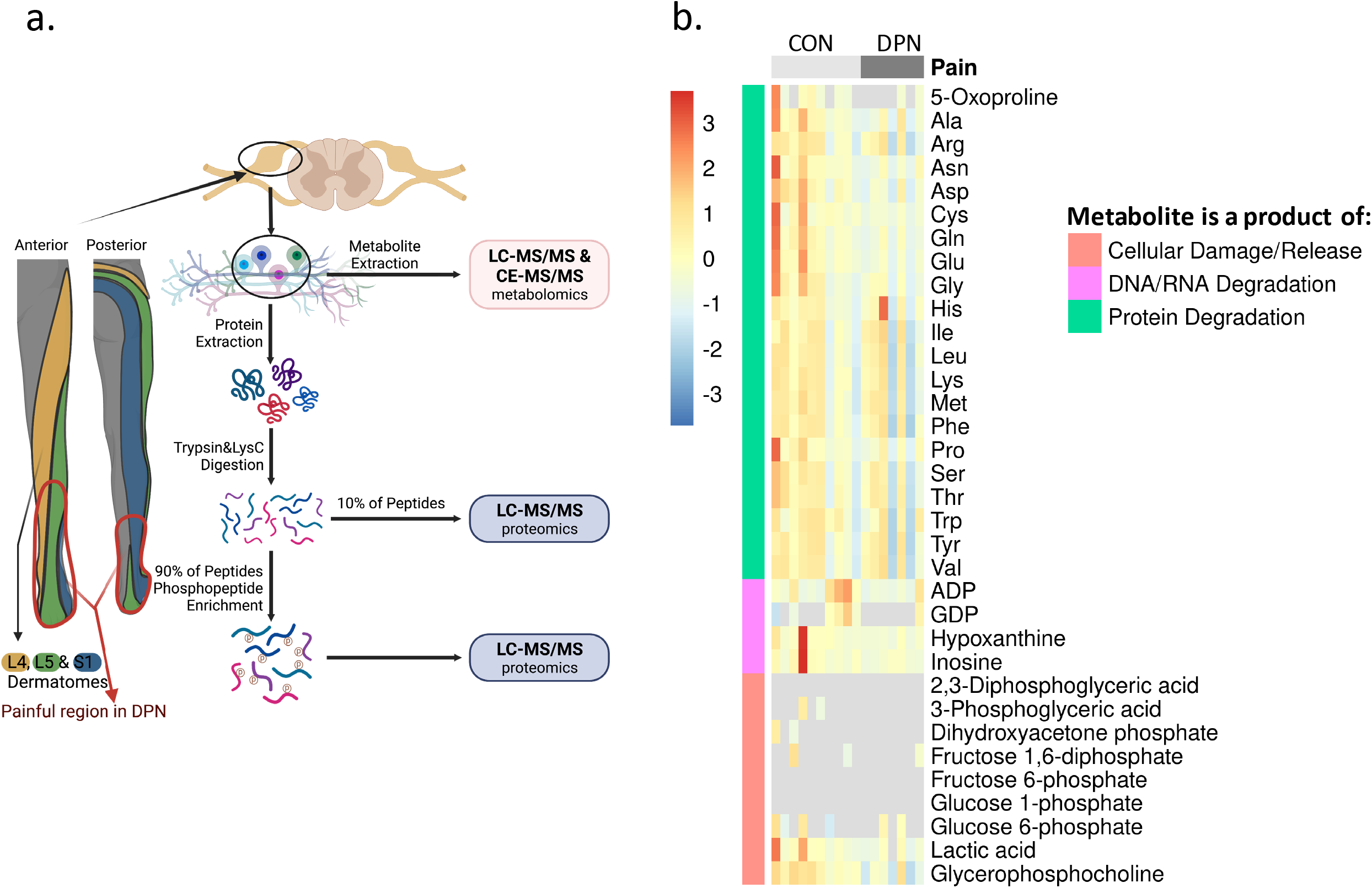
**A)** Schematic representation of study design. Innervation pattern for L4, L5, and S1 DRG used in analyses overlap with painful region in DPN. Created with BioRender. B) Heat map showing scaled and centered relative abundance data for select metabolites. Abnormally high profiles of all metabolites here would be indicative of tissue degradation. Two samples show higher abundance of some metabolites. However, these levels are reasonable for normal biological variation, and profiles of all noted metabolites are not observed to be consistently high. As such, DRG samples were determined to be of suitable quality for analysis. Row-wise annotations indicate degradative process which might produce indicated metabolites. CON indicates control samples, DPN indicates diabetic peripheral neuropathy.

**Supplementary Figure S2.**
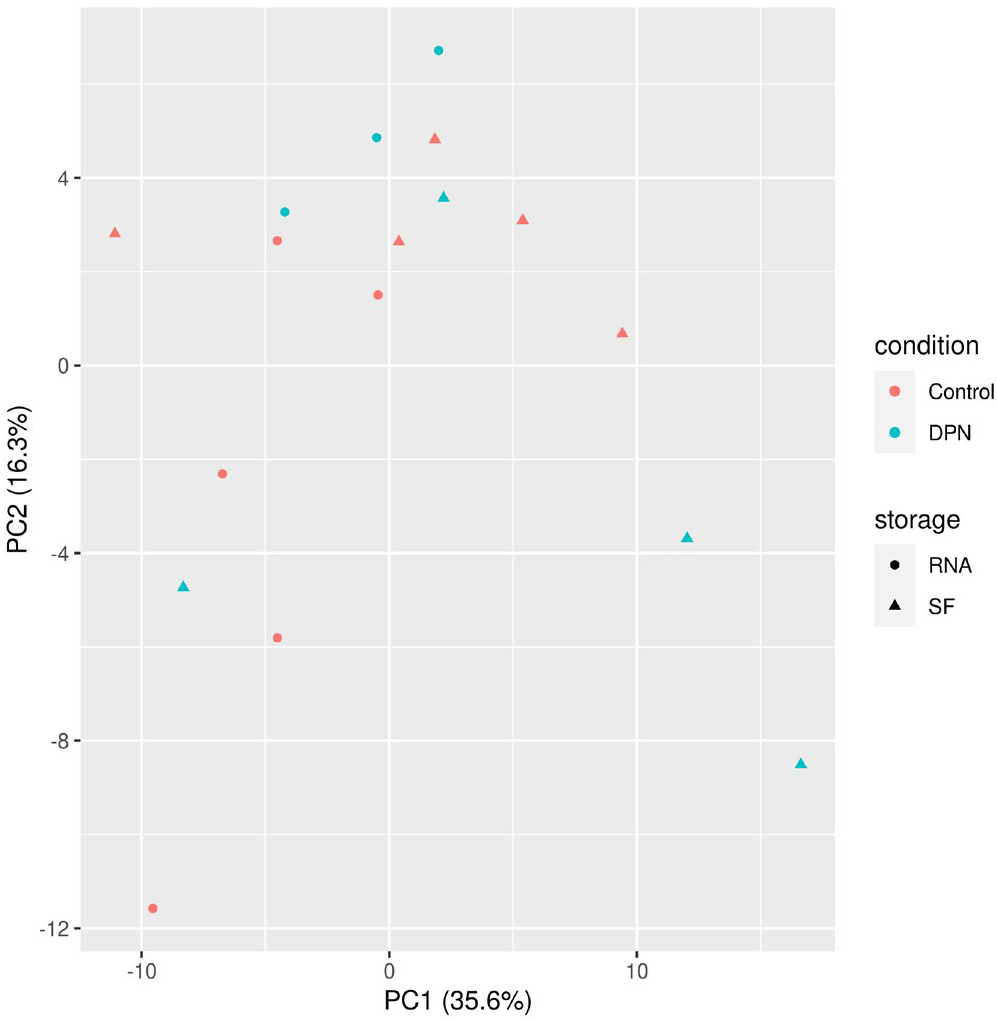
Plotted results of PCA performed with metabolomic data. DRG donor condition is indicated by color and tissue storage method (snap frozen (SF) or RNAlater (RNA)) is indicated by shape of the point.

**Supplementary Figure S3.**
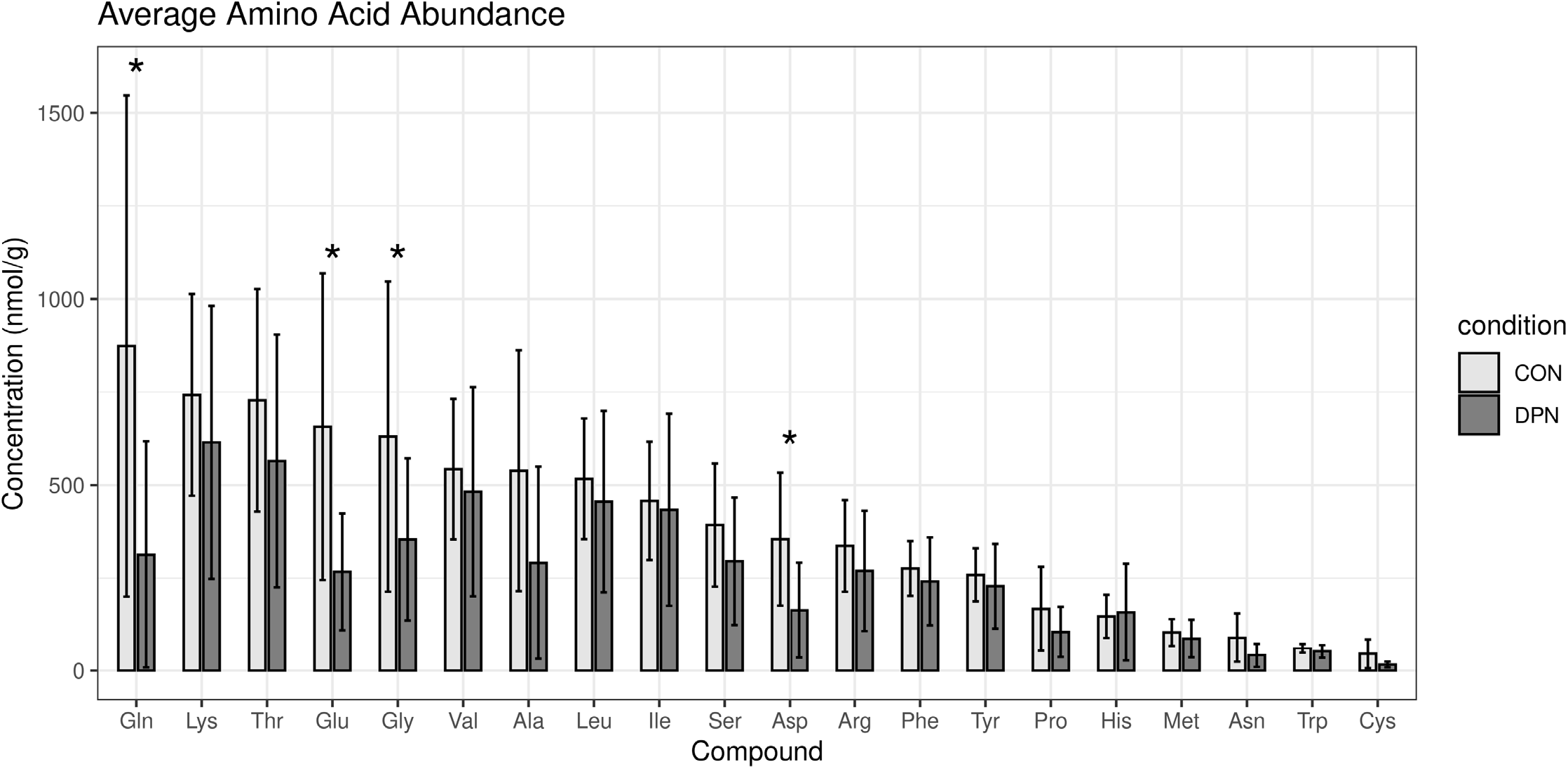
Concentration of amino acids in the DRG as detected using quantitative mass spectrometry panels. Amino acids are arranged according to highest average concentration in control human DRG tissues. CON indicates control, DPN indicates Diabetic Peripheral Neuropathy. Amino acids labeled with standard 3 letter abbreviations.

**Supplementary Figure S4.**
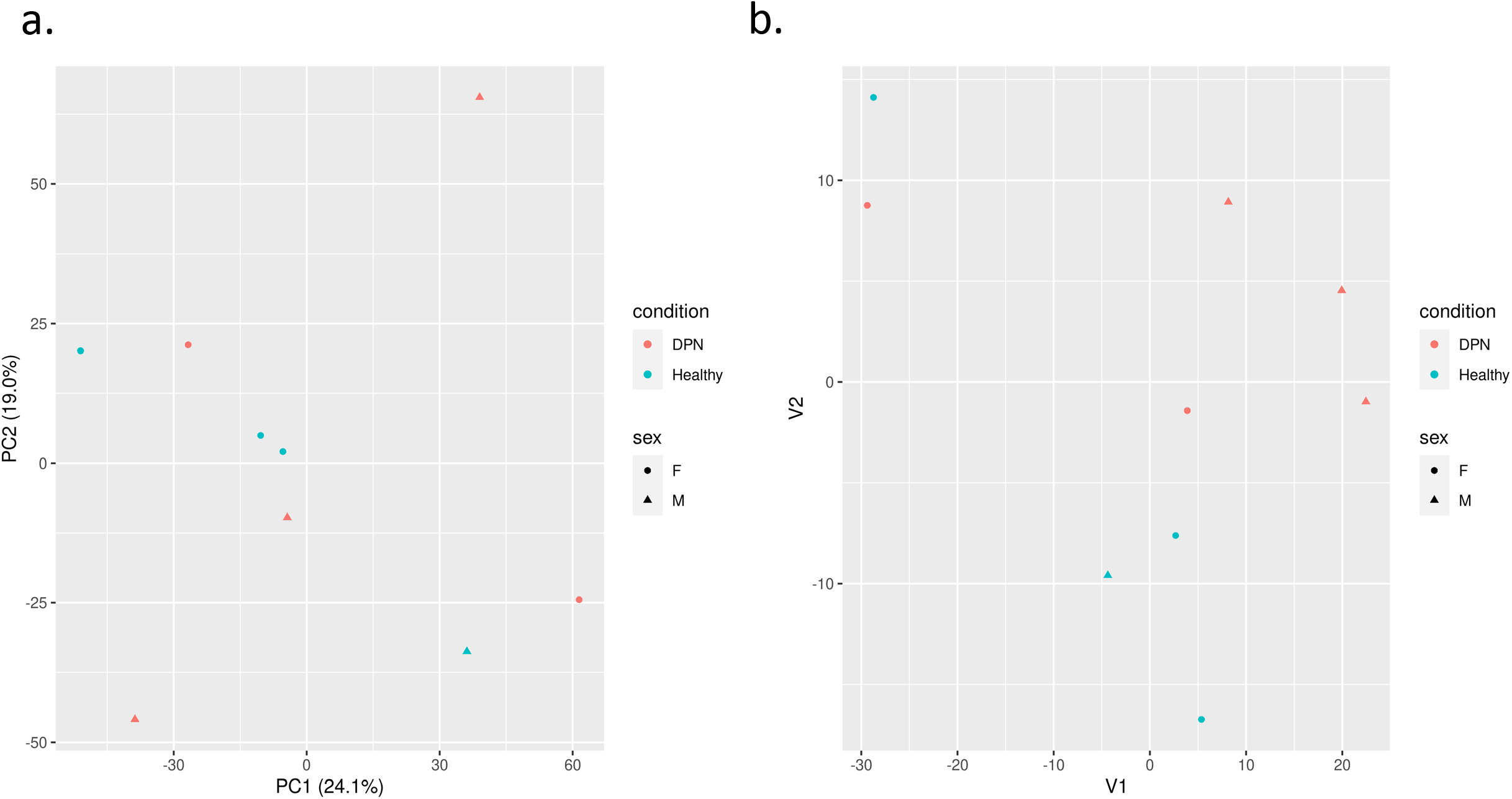
Plotted results of (A) PCA and (B) t-SNE performed with proteomic data. For both plots, DRG donor condition is indicated by color and donor sex (female (F) or male (M)) is indicated by shape of the point.

**Supplementary Figure S5.**
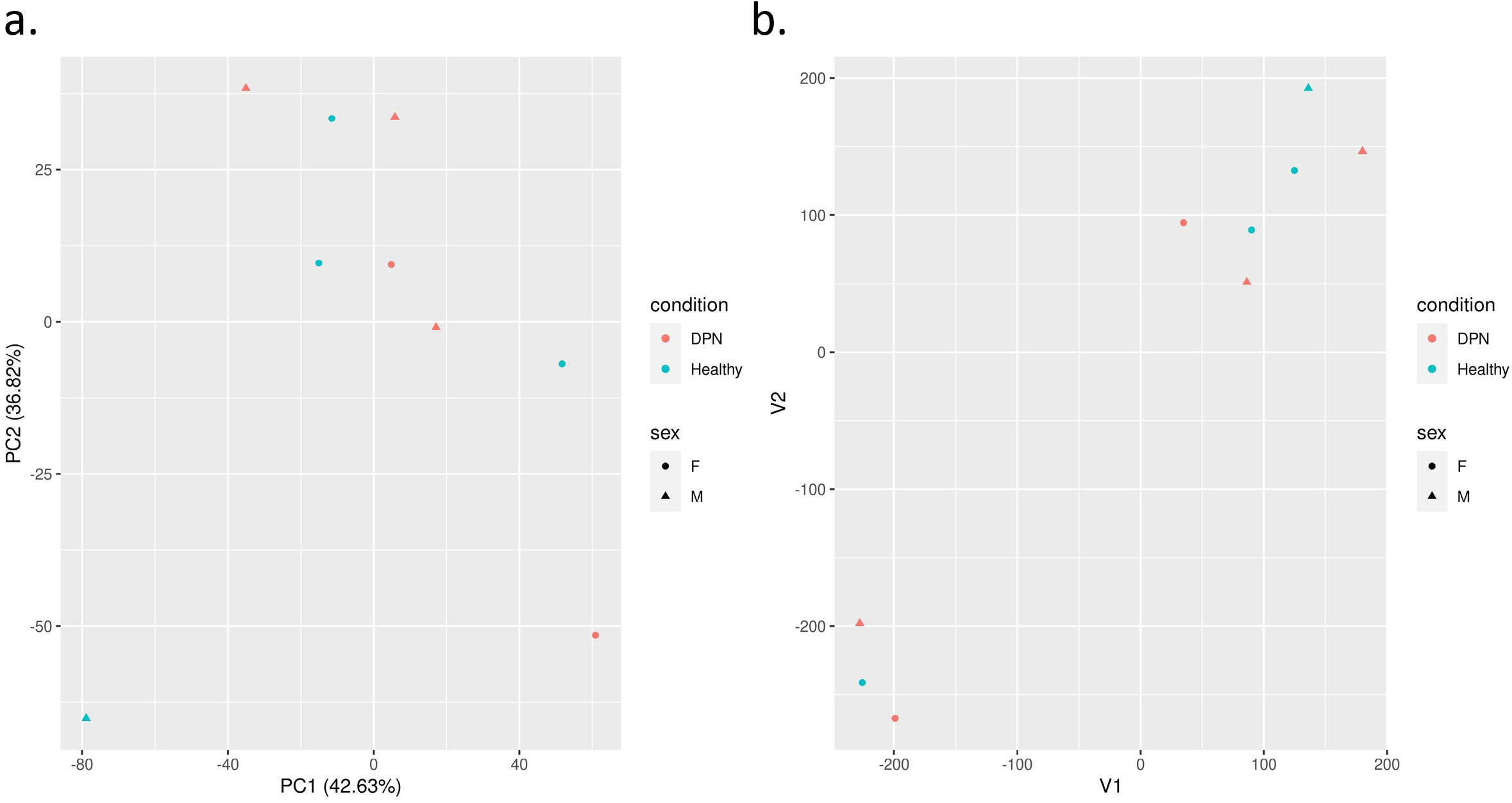
Plotted results of (A) PCA and (B) t-SNE performed with phospho-proteomic data. For both plots, DRG donor condition is indicated by color and donor sex (female (F) or male (M)) is indicated by shape of the point.

**Supplementary Figure S6.**
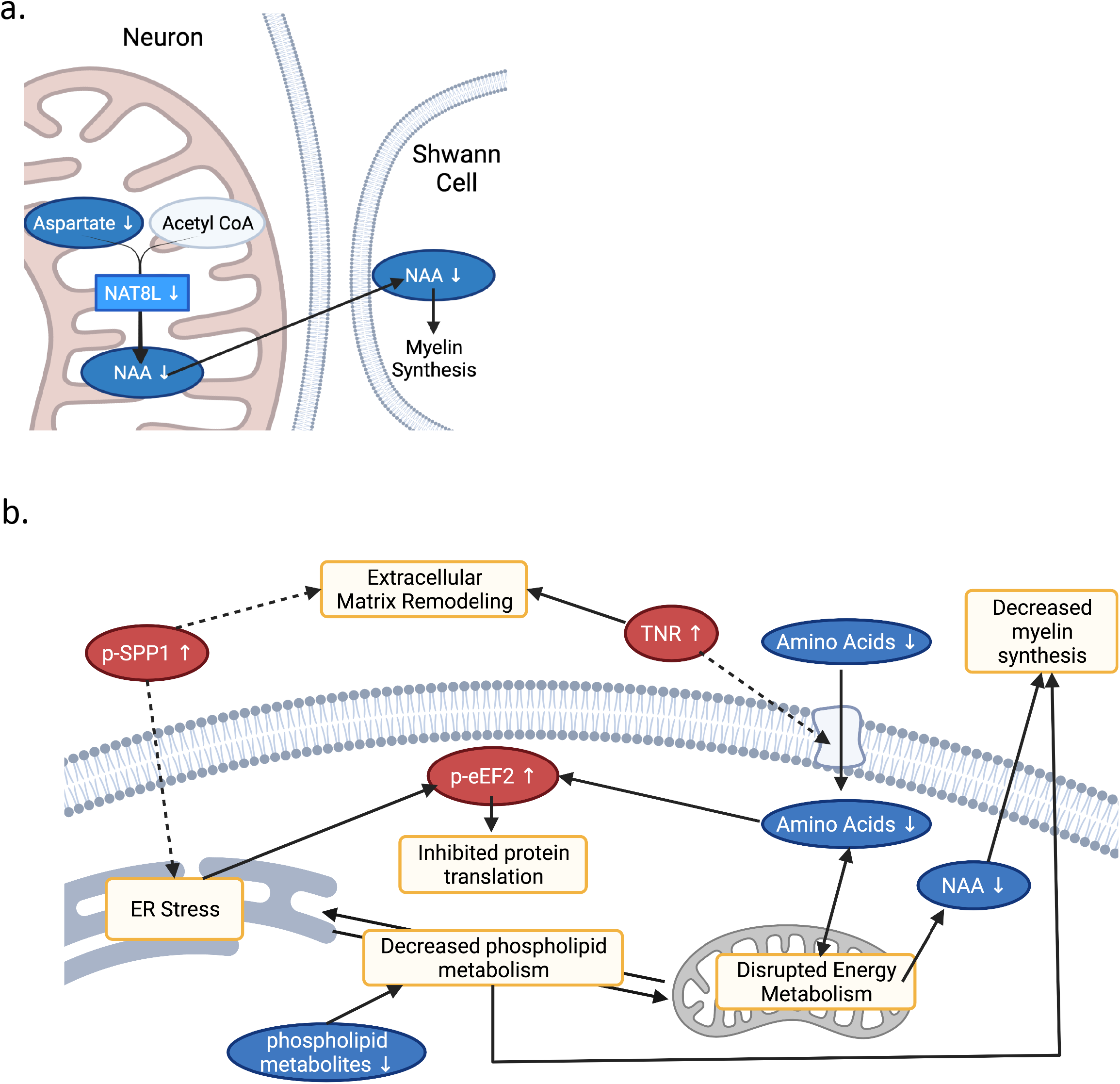
A) Schematic depicting a direct corelation between decreased metabolite NAA and decreased gene transcript NAT8L (created with BioRender). NAA is synthesized in the mitochondria from Asp by NAT8L. NAA is then transported to Schwann cells, where it is used as a precursor for myelin lipids. B) Schematic showing suggested interplay between identified molecular changes (created with BioRender). Briefly, amino acid starvation can impair mitochondrial function as well as inhibit protein translation, resulting in ER stress. Impaired function of the mitochondria and ER can also lead to disrupted crosstalk between these organelles and thus disrupted phospholipid synthesis. Disrupted mitochondrial function can also impair NAA synthesis, impacting glial-neuronal structures. Changes to the extracellular matrix can be related to ER stress and amino acid uptake. Finally, both ER stress and amino acid depletion can induce hyperphosphorylation of eEF2, thus inhibiting protein translation.

## References

1. Yong, R.J., P.M. Mullins, and N. Bhattacharyya, Prevalence of chronic pain among adults in the United States. Pain, 2022. 163(2): p. e328–e332.

2. Bouhassira, D., et al., Prevalence of chronic pain with neuropathic characteristics in the general population. Pain, 2008. 136(3): p. 380–387.

3. Khan, M.A.B., et al., Epidemiology of Type 2 Diabetes - Global Burden of Disease and Forecasted Trends. J Epidemiol Glob Health, 2020. 10(1): p. 107–111.

4. Saeedi, P., et al., Global and regional diabetes prevalence estimates for 2019 and projections for 2030 and 2045: Results from the International Diabetes Federation Diabetes Atlas, 9(th) edition. Diabetes Res Clin Pract, 2019. 157: p. 107843.

5. Tesfaye, S., et al., Diabetic neuropathies: update on definitions, diagnostic criteria, estimation of severity, and treatments. Diabetes Care, 2010. 33(10): p. 2285–93.

6. O’Brien, P.D., et al., Integrated lipidomic and transcriptomic analyses identify altered nerve triglycerides in mouse models of prediabetes and type 2 diabetes. Dis Model Mech, 2020. 13(2).

7. Smith, A.G. and J.R. Singleton, Obesity and hyperlipidemia are risk factors for early diabetic neuropathy. J Diabetes Complications, 2013. 27(5): p. 436–42.

8. Feldman, E.L., et al., New Horizons in Diabetic Neuropathy: Mechanisms, Bioenergetics, and Pain. Neuron, 2017. 93(6): p. 1296–1313.

9. Gregg, E.W., et al., Prevalence of lower-extremity disease in the US adult population >=40 years of age with and without diabetes: 1999-2000 national health and nutrition examination survey. Diabetes Care, 2004. 27(7): p. 1591–7.

10. Pop-Busui, R., et al., Diabetic Neuropathy: A Position Statement by the American Diabetes Association. Diabetes Care, 2017. 40(1): p. 136–154.

11. Krames, E.S., The dorsal root ganglion in chronic pain and as a target for neuromodulation: a review. Neuromodulation, 2015. 18(1): p. 24–32; discussion 32.

12. Hall, B.E., et al., Transcriptomic analysis of human sensory neurons in painful diabetic neuropathy reveals inflammation and neuronal loss. Sci Rep, 2022. 12(1): p. 4729.

13. Anderson, L. and J. Seilhamer, A comparison of selected mRNA and protein abundances in human liver. Electrophoresis, 1997. 18(3-4): p. 533–7.

14. Schwanhäusser, B., et al., Corrigendum: Global quantification of mammalian gene expression control. Nature, 2013. 495(7439): p. 126–7.

15. de Sousa Abreu, R., et al., Global signatures of protein and mRNA expression levels. Mol Biosyst, 2009. 5(12): p. 1512–26.

16. Schwaid, A.G., et al., Comparison of the Rat and Human Dorsal Root Ganglion Proteome. Sci Rep, 2018. 8(1): p. 13469.

17. Vogel, C. and E.M. Marcotte, Insights into the regulation of protein abundance from proteomic and transcriptomic analyses. Nat Rev Genet, 2012. 13(4): p. 227–32.

18. Zhang, S., et al., Insights Into Translatomics in the Nervous System. Front Genet, 2020. 11: p. 599548.

19. Weston, A.D. and L. Hood, Systems biology, proteomics, and the future of health care: toward predictive, preventative, and personalized medicine. J Proteome Res, 2004. 3(2): p. 179–96.

20. Freeman, O.J., et al., Metabolic Dysfunction Is Restricted to the Sciatic Nerve in Experimental Diabetic Neuropathy. Diabetes, 2016. 65(1): p. 228–38.

21. Hinder, L.M., et al., Decreased glycolytic and tricarboxylic acid cycle intermediates coincide with peripheral nervous system oxidative stress in a murine model of type 2 diabetes. J Endocrinol, 2013. 216(1): p. 1–11.

22. George, D.S., et al., Mitochondrial calcium uniporter deletion prevents painful diabetic neuropathy by restoring mitochondrial morphology and dynamics. Pain, 2022. 163(3): p. 560–578.

23. Leal-Julià, M., et al., Proteomic quantitative study of dorsal root ganglia and sciatic nerve in type 2 diabetic mice. Mol Metab, 2022. 55: p. 101408.

24. Percie du Sert, N. and A.S. Rice, Improving the translation of analgesic drugs to the clinic: animal models of neuropathic pain. Br J Pharmacol, 2014. 171(12): p. 2951–63.

25. Zisaki, A., L. Miskovic, and V. Hatzimanikatis, Antihypertensive drugs metabolism: an update to pharmacokinetic profiles and computational approaches. Curr Pharm Des, 2015. 21(6): p. 806–22.

26. Iadarola, M.J., M.R. Sapio, and A.J. Mannes, Be in it for the long haul: a commentary on human tissue recovery initiatives. J Pain, 2022.

27. Moffett, J.R., et al., N-Acetylaspartate reductions in brain injury: impact on post-injury neuroenergetics, lipid synthesis, and protein acetylation. Front Neuroenergetics, 2013. 5: p. 11.

28. Ryazanov, A.G., Ca2+/calmodulin-dependent phosphorylation of elongation factor 2. FEBS Lett, 1987. 214(2): p. 331–4.

29. Carlberg, U., A. Nilsson, and O. Nygård, Functional properties of phosphorylated elongation factor 2. Eur J Biochem, 1990. 191(3): p. 639–45.

30. Gauthier-Coles, G., et al., Quantitative modelling of amino acid transport and homeostasis in mammalian cells. Nat Commun, 2021. 12(1): p. 5282.

31. Adeva-Andany, M., et al., Insulin resistance and glycine metabolism in humans. Amino Acids, 2018. 50(1): p. 11–27.

32. Menge, B.A., et al., Selective amino acid deficiency in patients with impaired glucose tolerance and type 2 diabetes. Regul Pept, 2010. 160(1-3): p. 75–80.

33. Sadasivan, S., et al., Amino acid starvation induced autophagic cell death in PC-12 cells: evidence for activation of caspase-3 but not calpain-1. Apoptosis, 2006. 11(9): p. 1573–82.

34. Sato, H., et al., Neurodegenerative processes accelerated by protein malnutrition and decelerated by essential amino acids in a tauopathy mouse model. Sci Adv, 2021. 7(43): p. eabd5046.

35. Suraweera, A., et al., Failure of amino acid homeostasis causes cell death following proteasome inhibition. Mol Cell, 2012. 48(2): p. 242–53.

36. Wiame, E., et al., Molecular identification of aspartate N-acetyltransferase and its mutation in hypoacetylaspartia. Biochem J, 2009. 425(1): p. 127–36.

37. Mersmann, N., et al., Aspartoacylase-lacZ knockin mice: an engineered model of Canavan disease. PLoS One, 2011. 6(5): p. e20336.

38. Zaroff, S., et al., Transcriptional regulation of N-acetylaspartate metabolism in the 5xFAD model of Alzheimer’s disease: evidence for neuron-glia communication during energetic crisis. Mol Cell Neurosci, 2015. 65: p. 143–52.

39. Maccarrone, M., et al., Cannabimimetic activity, binding, and degradation of stearoylethanolamide within the mouse central nervous system. Mol Cell Neurosci, 2002. 21(1): p. 126–40.

40. Osman, C., D.R. Voelker, and T. Langer, Making heads or tails of phospholipids in mitochondria. J Cell Biol, 2011. 192(1): p. 7–16.

41. Poitelon, Y., A.M. Kopec, and S. Belin, Myelin Fat Facts: An Overview of Lipids and Fatty Acid Metabolism. Cells, 2020. 9(4).

42. Bogen, O., et al., GDNF hyperalgesia is mediated by PLCgamma, MAPK/ERK, PI3K, CDK5 and Src family kinase signaling and dependent on the IB4-binding protein versican. Eur J Neurosci, 2008. 28(1): p. 12–9.

43. Li, Z.Z., et al., Extracellular matrix protein laminin beta1 regulates pain sensitivity and anxiodepression-like behaviors in mice. J Clin Invest, 2021. 131(15).

44. Reichling, D.B., P.G. Green, and J.D. Levine, The fundamental unit of pain is the cell. Pain, 2013. 154 Suppl 1: p. S2–9.

45. Okuda, H., et al., Chondroitin sulfate proteoglycan tenascin-R regulates glutamate uptake by adult brain astrocytes. J Biol Chem, 2014. 289(5): p. 2620–31.

46. Reinhard, J., L. Roll, and A. Faissner, Tenascins in Retinal and Optic Nerve Neurodegeneration. Front Integr Neurosci, 2017. 11: p. 30.

47. Boyce, M., et al., A pharmacoproteomic approach implicates eukaryotic elongation factor 2 kinase in ER stress-induced cell death. Cell Death Differ, 2008. 15(3): p. 589–99.

48. Henry, S.A., S.D. Kohlwein, and G.M. Carman, Metabolism and regulation of glycerolipids in the yeast Saccharomyces cerevisiae. Genetics, 2012. 190(2): p. 317–49.

49. Barragan-Iglesias, P., et al., Activation of the integrated stress response in nociceptors drives methylglyoxal-induced pain. Pain, 2019. 160(1): p. 160–171.

50. Inceoglu, B., et al., Endoplasmic reticulum stress in the peripheral nervous system is a significant driver of neuropathic pain. Proc Natl Acad Sci U S A, 2015. 112(29): p. 9082–7.

51. Lindholm, D., H. Wootz, and L. Korhonen, ER stress and neurodegenerative diseases. Cell Death Differ, 2006. 13(3): p. 385–92.

52. Öztürk, Z., C.J. O’Kane, and J.J. Pérez-Moreno, Axonal Endoplasmic Reticulum Dynamics and Its Roles in Neurodegeneration. Front Neurosci, 2020. 14: p. 48.

53. Lin, W. and B. Popko, Endoplasmic reticulum stress in disorders of myelinating cells. Nat Neurosci, 2009. 12(4): p. 379–85.

54. Goswami, Samridhi C et al. Molecular signatures of mouse TRPV1-lineage neurons revealed by RNA-Seq transcriptome analysis. The journal of pain, 2014. 15(12): p. 1338–1359.

55. Marsh BC, Kerr NC, Isles N, Denhardt DT, Wynick D. Osteopontin expression and function within the dorsal root ganglion. Neuroreport, 2007. 18(2): p. 153–7.

56. Dalal, S., et al., Osteopontin stimulates apoptosis in adult cardiac myocytes via the involvement of CD44 receptors, mitochondrial death pathway, and endoplasmic reticulum stress. Am J Physiol Heart Circ Physiol, 2014. 306(8): p. H1182–91.

57. Wu, J., et al., NK cells induce hepatic ER stress to promote insulin resistance in obesity through osteopontin production. J Leukoc Biol, 2020. 107(4): p. 589–596.

58. Chai, Y.L., et al., Plasma osteopontin as a biomarker of Alzheimer’s disease and vascular cognitive impairment. Sci Rep, 2021. 11(1): p. 4010.

59. Kahles, F., H.M. Findeisen, and D. Bruemmer, Osteopontin: A novel regulator at the cross roads of inflammation, obesity and diabetes. Mol Metab, 2014. 3(4): p. 384–93.

60. Giachelli, C.M. and S. Steitz, Osteopontin: a versatile regulator of inflammation and biomineralization. Matrix Biol, 2000. 19(7): p. 615–22.

61. Jakovcevski, I., et al., Tenascins and inflammation in disorders of the nervous system. Amino Acids, 2013. 44(4): p. 1115–27.

62. Rowe, G.C., et al., PGC-1alpha induces SPP1 to activate macrophages and orchestrate functional angiogenesis in skeletal muscle. Circ Res, 2014. 115(5): p. 504–17.

63. M, S.L. and P. O, Inflammatory biomarkers as a part of diagnosis in diabetic peripheral neuropathy. J Diabetes Metab Disord, 2021. 20(1): p. 869–882.

64. Tsalamandris, S., et al., The Role of Inflammation in Diabetes: Current Concepts and Future Perspectives. Eur Cardiol, 2019. 14(1): p. 50–59.

65. Wei, R., et al., Missing Value Imputation Approach for Mass Spectrometry-based Metabolomics Data. Sci Rep, 2018. 8(1): p. 663.

66. Kendrick, N., et al., Preparation of a phosphotyrosine-protein standard for use in semiquantitative western blotting with enhanced chemiluminescence. PLoS One, 2020. 15(6): p. e0234645.

